# Thioflavin T In-gel Stain to Study Protein Misfolding in Frozen Tissue Specimens

**DOI:** 10.1101/2023.05.12.540528

**Authors:** Joseph Oldam, Irina Tchernyshyov, Jennifer E. Van Eyk, Juan Troncoso, Charles G. Glabe, Giulio Agnetti

## Abstract

There are limited options to quantify and characterize amyloid species from biological samples in a simple fashion. Thioflavin T (ThT) has now been used for decades to stain amyloid fibrils but to our knowledge we were the first to use it in-gel. Thioflavin T in-gel stain is convenient as it is fast, inexpensive, available to most labs, compatible with other fluorescent stains and downstream analyses such as mass spectrometry (MS).

## Main

Organ and systemic proteinopathies represent one of the main area of public health concern in Westernized society in this century^1^. The toxic nature of pre-fibrillar and fibrillar amyloid is well-established in neurological disease^2^, however their role and prevalence are emerging in cardiovascular disease as well^3^. Most amyloid species are quantitated by either fluorescent, radiolabeled, or immunological probes as well as by mass spectrometry (MS)^4-7^. From the technological standpoint, MS is arguably one of the techniques with the highest impact on biomedical sciences in this century^8^. Cutting-edge MS-based approaches such as isotope incorporation are currently being used in the study of protein misfolding but their application is still confined to few laboratories worldwide^9^. It is our opinion that a simple way to detect and quickly isolate amyloid species *ex vivo*, which is compatible with “classic” MS, would greatly benefit the study of an increasing number of organ-based proteinopathies.

We optimized a straightforward, affordable way to stain amyloid species in-gel, and tested it on a well-established model of murine cardiac amyloidosis^10^, a kind gift of Dr. Jeffrey Robbins and colleagues. The idea of an in-gel fluorescent staining for amyloids came about after reading the ingenious study from Herve and colleagues who utilized Thioflavin T (ThT) in combination with a general, fluorescent protein stain (SYPRO Ruby) to assess cross-contamination of surgical tools with prions^11^. As SYPRO Ruby is typically used to stain proteins separated by gel electrophoresis we hypothesized that ThT would work in-gel as well.

We combined the staining with both a modified version of blue-native gels in the presence of SDS, which we referred to as Not-So-Native (NSN) as well as regular SDS-PAGE. These approaches have the advantage of separating oligomers and fibrils, combined with the ease of quantifying these species in a simple gel format and on a wide range of molecular sizes. In fact, ThT and other related compounds have an affinity for both fibrils and pre-fibrillar oligomers^12,13^ (named pre-amyloid oligomers, or PAOs henceforth) and its fluorescence increases when ThT is bound to them^14^.

Thioflavin T’s signal can be conveniently acquired using a laser scanner equipped with a Cy2 filter (λ_ex/em_=488/520). A Cy5 (λ_ex/em_=633/670) filter can be used as a reference filter and for total protein quantitation based on Coomassie staining. In short, these aspects combined with the low cost of the dye make the described method accessible to a variety of laboratories. The additional optimization of a classical reducing method also allows us to compare the results obtained using ThT with those obtained using more classical approaches (e.g. the A11 antibody^15^). For these reasons, we believe that this new approach will enable the expeditious quantification, purification and characterization of amyloid species in different proteinopathies (e.g. cardiac amyloidosis, Alzheimer’s, Parkinson’s, etc.).

We first used the ThT in-gel stain to validate the amyloid properties of desmin oligomers in a canine model of dyssynchronous heart failure (HF)^16^. In the study we separated the oligomers under non-reducing conditions and in the presence of sodium dodecyl sulphate (SDS), using a blue-native gel format, which we nicknamed not-so-native (NSN).

More recently, we tested the performance of the staining and optimized the protocol for classical (reducing) SDS-PAGE using the best characterized model of cardiac proteotoxicity^10,17^. The transgenic mice used in this study express a mutated form (R120G) of the most abundant small heat shock protein in the heart: α-B-crystallin (cryAB). The cardiac-specific expression of this mutated chaperone in transgenic mice induces the formation of desmin PAOs and fibrils.

In the optimized protocol outlined in **Figure 1**, cardiac extracts are prepared according to our optimized protocol to separate myofilament and cytosolic-enriched fractions^17^. This separation consists of an homogenization step in the absence of detergents (cytosolic-enriched, soluble fraction), followed by centrifugation and re-suspension in the presence of detergents (SDS for the myofilament-enriched, insoluble fraction), similar to the protocols used in neuroscience to enrich for fibrils^18^ (**Figure 1a**). After protein quantification (**Figure 1b**), samples are diluted in the proper sample buffer for either NSN- or classical SDS-PAGE, boiled in the latter case and separated using a standard 1D-PAGE apparatus (**Figure 1c**). After separation, the gel, which is already blue in the case of NSN and clear in the case of regular denaturing gels, is fixed, rinsed thoroughly in water, stained with Coomassie for the reducing gels, and its images are digitized prior to staining with ThT to ensure that potential protein auto-fluorescence in the ThT (Cy2) channel is accounted for (**Figure 1d**). The gel is then incubated with 50 *μ*M ThT in acidified water, rinsed several times in acidified water to reduce background and remove ThT speckles, prior to the final image acquisition (**Figure 1d**, see Methods for details). Following image acquisition, the gel can be stored in bi-distilled water or dried. The ThT-positive gel bands can be located using the Coomassie image as a reference and excised for downstream in-gel digestion and MS analysis (**Figures 1e and 1f**).

**Figure 1.**
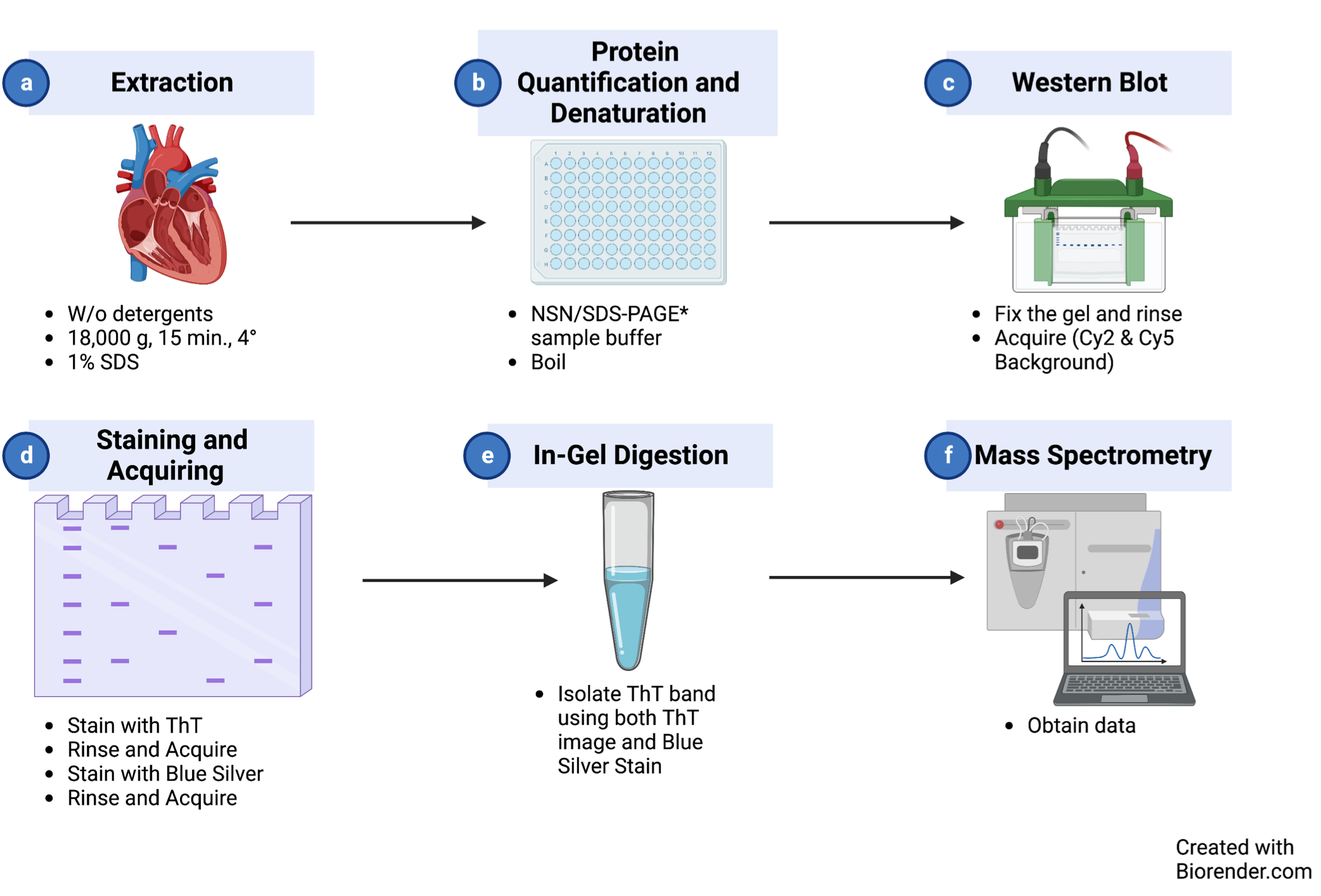
Thioflavin T Staining/MS Workflow.

When this procedure was used to analyze short cardiac amyloid fibrils by NSN-PAGE, we could detect a ∼5-fold increase of a sharp ThT-positive band in R120G cryAB (R120G) mice *vs*. non-transgenic (NTG) controls (*P*=0.0002). Signals for both ThT and Coomassie were digitized and contrast-enhanced (**Figure 2a**) to enable improved visualization and quantitation (**Figure 2b**). Interestingly the ThT-positive band was detected at the apparent molecular weight of ∼600 kDa. Although NSN separation does not allow for a proper molecular weight calibration, we reported a similar electrophoretic mobility of ThT-positive bands in canine failing hearts^16^, suggesting a mechanism of fibrillization which is conserved across species and independent from genetic mutations. This is relevant as many organ proteinopathies (e.g. Parkinson) are highly sporadic in nature^3^.

**Figure 2.**
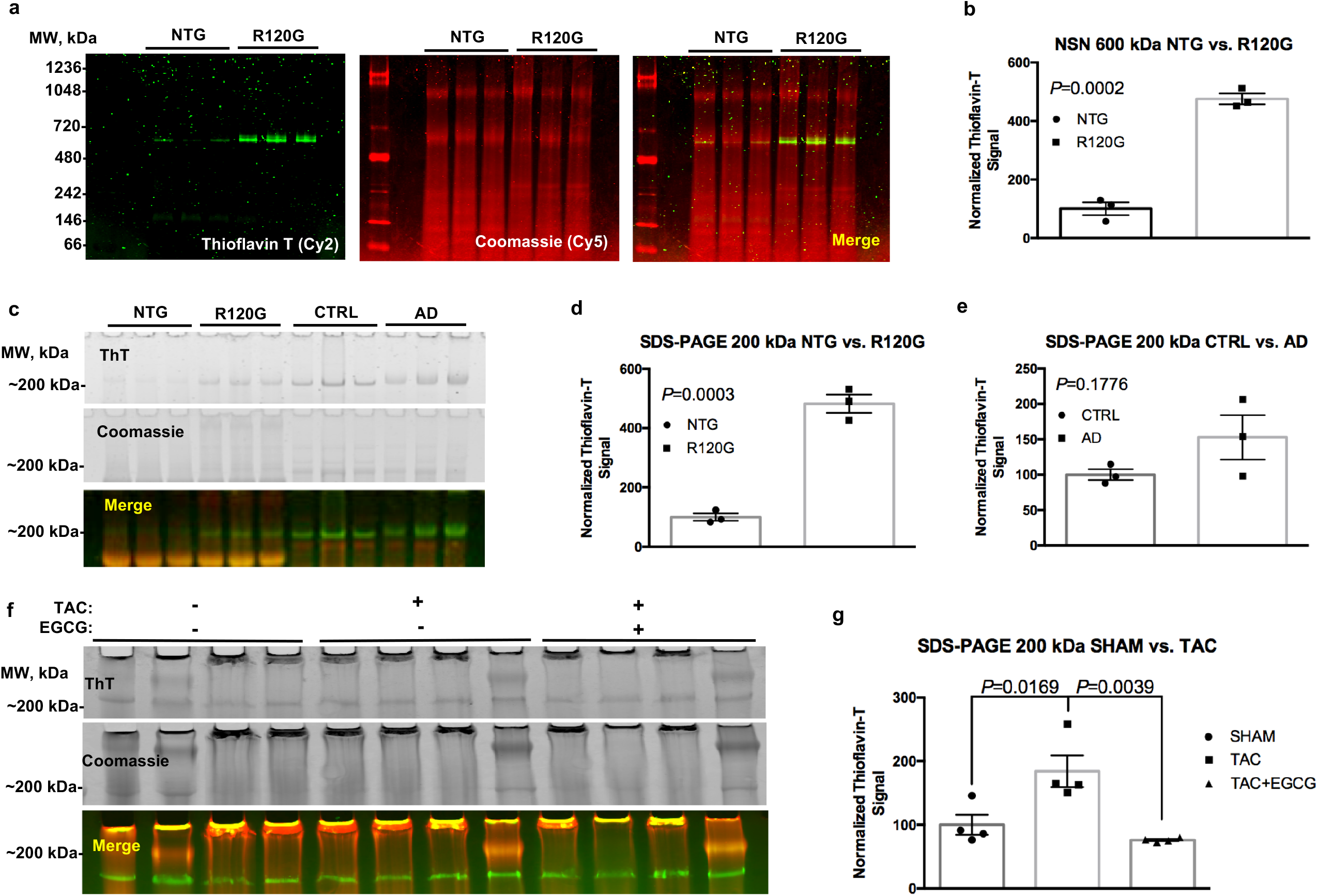
ThT Stain in Cardiac and Brain Tissue Extracts. Representative image of a not-so-native (NSN) gel used to separate fibrillar aggregates from NTG and R120G cryAB mice (**a**, ThT in green, Coomassie in red); corresponding densitometric analysis for the ThT signal at ∼600 kDa is also provided (**b**). Representative image of a classical SDS-PAGE gel showing a ThT signal at ∼200 kDa in NTG and R120G cryAB mouse cardiac extracts as well as extracts from healthy human brain (CTRL) and Alzheimer’s (AD) human brain tissues. Densitometric analysis for the ThT signal at ∼200 kDA is also provided (**d** and **e**, respectively). A representative image of the effects of EGCG on extracts from TAC and sham-operated mouse hearts is also provided (**f**), along with the respective densitometric analysis (**g**). NTG, non-transgenic; R120G, R120G cryAB mouse heart extracts; CTRL, healthy human brain tissue; AD, Alzheimer disease human brain tissue; EGCG, epigallocatechin gallate; ThT, Thioflavin T. Mean ± SD is plotted in all graphs. *P*-values were obtained through student’s *t*-test (**b**,**d-e**) or one-way ANOVA followed by Sidak’s multiple comparison test (**g**). Please refer to the text for the abbreviations.

In order to widen the applicability of the protocol we adapted it to use reducing (classical) SDS-PAGE. We tested the method using both R120G cryAB samples *vs*. non-transgenic controls (NTG), and clinical brain specimens from Alzheimer’s (AD) patients *vs*. healthy controls (CTRL) (**Figure 2c**). In agreement with our previously published results and the NSN protocol, we detected a ∼5-fold significant increase in a ThT-positive band at ∼200 kDa under reducing conditions in cryAB R120G cardiac extracts vs. NTG controls (*P*=0.0003 **Figure 2d-e**). We also detected a ∼1.5-fold increase in a ∼200 kDa ThT-positive band compatible with fibrils in brain extracts from Alzheimer’s patients compared to healthy controls (*P*=0.0176 **Figure 2e**). We attribute the lack of significance with the latter comparison to the limited number of samples utilized in this study and the larger variability in patients’ cohorts compared to mice with homogeneous genetic background. As mentioned, Coomassie post-stain allowed us to cross-reference and to excise the ThT-positive bands with the naked eye (**Figure 2c**, mid panel), as well as confirming equal loading. In these settings we were able to confirm the specificity of the stain using a standardized and more robust separation method (classical SDS-PAGE).

Lastly, we used the ThT in-gel protocol to measure the accumulation of desmin-positive amyloid fibrils in an established murine model of HF based on transverse aortic constriction (TAC) vs. sham-operated controls)^17^. We detected a ∼2-fold increase in a ∼200 kDa ThT-positive band in extracts from TAC mice samples *vs*. controls (*P*=0.0169, **Figure 2g**).

In order to further validate the specificity of the ThT-staining for amyloid aggregates, we exploited the ability of the small molecule epigallocatechin gallate (EGCG) to reduce or reverse fibrillization in prion strains^19^. Of note, treatment cardiac extracts with 100 *μ*M EGCG for 30 min (RT) reversed the increase in the ThT observed with TAC samples to control levels (*P*=0.0039) (**Figure 2g**). This observation supports the specificity of the ThT-staining for amyloid species. We also obtained similar results on the effect of in vitro treatment with EGCG on extracts from human brains Alzheimer’s patients *vs*. healthy controls. These combined data suggest that the ThT in-gel staining could be applicable to a wide number of proteinopathies, including those afflicting organs other than the heart (e.g. the brain).

## Methods

### Tissue procurement

8-12 week old C57BL/6 mice were subjected to TAC or sham surgery through the Cardiac Physiology core at Johns Hopkins as previously described^17,20^. Four weeks after surgery TAC mice develop overt HF as measured by a drop in fractional shortening (≤40%) by echo. At that time point mice where anesthetized and the heart was harvested and snap frozen according to the protocol approved by the local IACUC. Briefly, mice where anesthetized using isoflurane. Upon reaching deep anesthesia with isoflurane (absence of toe pinch reflex), cervical vertebrae were dislocated, and the heart was quickly extracted through thoracotomy. Freshly explanted hearts were quickly rinsed in ice-cold PBS containing protease (cOmplete Mini, Roche) and phosphatase inhibitors (PhosStop, Sigma) to remove excess blood, gently blotted on clean filter paper to remove excess liquid, snap-frozen in liquid nitrogen and stored at -80C until further processing. Frozen ventricles from R120G cryAB mice were kindly provided by Dr. Jeff Robbins at the Cincinnati Children’s Hospital. Frozen brain tissue from Alzheimer’s patients and healthy controls (n=3 ea.), were the Johns Hopkins University Brain Resource Center and Neuropathology Core of the Alzheimer’s Disease Research Center. All frozen tissue samples were stored at -80 °C until ready for protein extraction.

### Protein Extraction

30-50 mg of frozen tissue was homogenized in cold PBS (Gibco), completed with protease (cOmplete Mini, Roche) and phosphatase (PhosStop, Sigma) inhibitors, and 25 mM Hepes (Amresco). Pre-weighed 2 ml vials were used to obtain the tissue weight. A dry ice-cooled metal bead (Retsch) was placed in each vial along with 5 volumes (V/W) of homogenization buffer. Samples where then homogenized for 2 min at 28 Hz using a bead mill (MM400, Retsch) and cooled racks. After milling, samples were pulse centrifuged, placed in a magnetic rack (Invitrogen) to ensure that the metal bead would stay on the side of the vial, and the homogenate transferred to a new clean vial on ice. The beads were rinsed with 3 volumes of ice-cold homogenization buffer (V/W) under vortexing and the resulting solution was reunited with the respective homogenate. The resulting tissue homogenates were centrifuged for 15 min (18,000 rcf, 4 °C) to obtain soluble, and insoluble fractions which were separated, snap-frozen and stored at -80ºC until further use.

### Protein Denaturation and Quantification

Soluble and insoluble fractions were denatured in LDS buffer (Thermo) in homogenization buffer, completed with 1% DTT (V/V) followed by heat denaturation at 95 °C for 10’. Protein concentration in the denatured samples was measured by the EZQ Protein Quantification Kit (Thermo).

### Electrophoresis

For not-so-native (NSN) gels, protein extracts from the insoluble fraction were diluted in blue native sample buffer (25 mM BisTris, Bio-Rad; 0.015 1N Hydrochloric Acid, Fisher Scientific; 10% glycerol, Sigma Aldrich; 25 mM NaCl, Sigma Aldrich; 0.001% Ponceau S, Fisher Scientific; 2% SDS, Fisher Scientific). After incubating samples at RT for 30 min, samples were clarified by centrifugation (18,000 rcf, 30 min, 4 °C) to remove the insoluble material and the resulting supernatant was completed with 0.25% Coomassie Blue G-250 (Fisher Scientific). Forty *μ*g of protein per sample were then separated using NuPAGE pre-cast native gels (Thermo) for NSN while 20 *μ*g of protein were used for classic SDS-PAGE using 3-12% NuPAGE gels (Thermo). Both type of electrophoreses were performed at 150 V for ∼1 hour. After separation, gels were fixed in 10% acetic acid (Fisher Chemical), 40% methanol (Fisher Chemical), in bi-distilled water overnight to remove excess of SDS for downstream excess analysis.

### Gel Staining/Acquisition

The following day gels were rinsed several washes in bi-distilled water to remove excess methanol, and the background fluorescent signal for ThT and Coomasie (Cy2 and Cy5 filters, respectively) using a Typhoon laser scanner (GE). Gels were subsequently stained with 0.1% (W/V) ThT in acidified water (13.8 *μ*l of 1N Hydrochloric Acid (Fisher Scientific) per 50 ml of bi-distilled water) for 1 hour in the dark. The ThT working solution was prepared at least one hour ahead of time to maximize solubilization and kept in the dark. After ThT-staining, gels were rinsed extensively in acidified water to remove speckles followed by a new acquisition of the Cy2 and Cy5 signal using the Typhoon. Following ThT-signal acquisition, gels were stained in blue silver^21^ for ≥ 20 min, RT rinsed extensively in acidified water and imaged again to acquire the fluorescent signal for the Coomassie (Cy5 filter). The resulting, merged image of Coomassie and ThT signals (**Figure 2**) was utilized for quantitative analysis as well as to locate the ThT-positive bands with the naked eye using Coomassie as a reference.

### Mass Spectrometry

please refer to the online Material, Methods and Results for LC-MS analysis (**Supplementary Figure 1-2**).

## Supporting information

Supplemental Methods and Results

## Contributions

JO replicated experiments and contributed to the writing of the experimental portions of the manuscript. IC and JE performed the MS analysis. JT provided access to brain tissue form patients while CG provided critical feedback for the development of the method. GA conceived and optimized the method and wrote the manuscript.

## Conflict of Interest

The Authors have no conflict of interest to disclose.

## Acknowledgments

The Authors are grateful to the Johns Hopkins University Alzheimer’s Disease Research Center (NIH P30AG066507) for providing access to patient tissue specimens.

## Source of Funding

The study was supported by the Zegar Family Foundation, 18TPA34170382 from the American Heart Association, the Magic that Matters Foundation and by the Leducq Foundation TNE ID#673168 (to GA).

## Notes

https://www.agnettilab.com

### Competing Interest Statement

The authors have declared no competing interest.

