## Supplementary Figures for "Thioflavin T In-gel Stain to Study Protein Misfolding in Frozen Tissue Specimens"

### Supplementary Figure 1

**a**

DESM_MOUSE (100%), 53,498.7 Da
Desmin OS=Mus musculus GN=Des PE=1 SV=3
8 exclusive unique peptides, 8 exclusive unique spectra, 16 total spectra, 100/469 amino acids (21% coverage)

```
MSQAYSSSQR VSSYRRTFGG APGFSLGSPL SSPVFPRAGF GTKGSSSSMT SRVYQVSRTS GGAGGLGSLR
SSRLGTTRAP SYGAGELLDF SLADAVNQEF LATRTNEKVE LQELNDRFAN YIEKVRFLEQ QNAALAAEVN
RLKGREPTRV AELYEEEMRE LRRQVEVLTN QRARVDVERD NLIDDLQRLK AKLQEEIQLR EEAENNLAAF
RADVDAATLA RIDLERRIES LNEEIAFLKK VHEEEIRELQ AQLQEQQVQV EMDMSKPDLT AALRDIRAQY
ETIAAKNISE AEEWYKSKVS DLTQAANKNN DALRQAKQEM MEYRHQIQSY TCEIDALKGT NDSLMRQMRE
LEDRFASEAN GYQDNIARLE EEIRHLKDEM ARHLREYQDL LNVKMALDVE IATYRKLLEG EESRINLPIQ
TFSALNFRET SPEQRGSEVH TKKTVMIKTI ETRDGEVVSE ATQQQHEVL
```

**b**

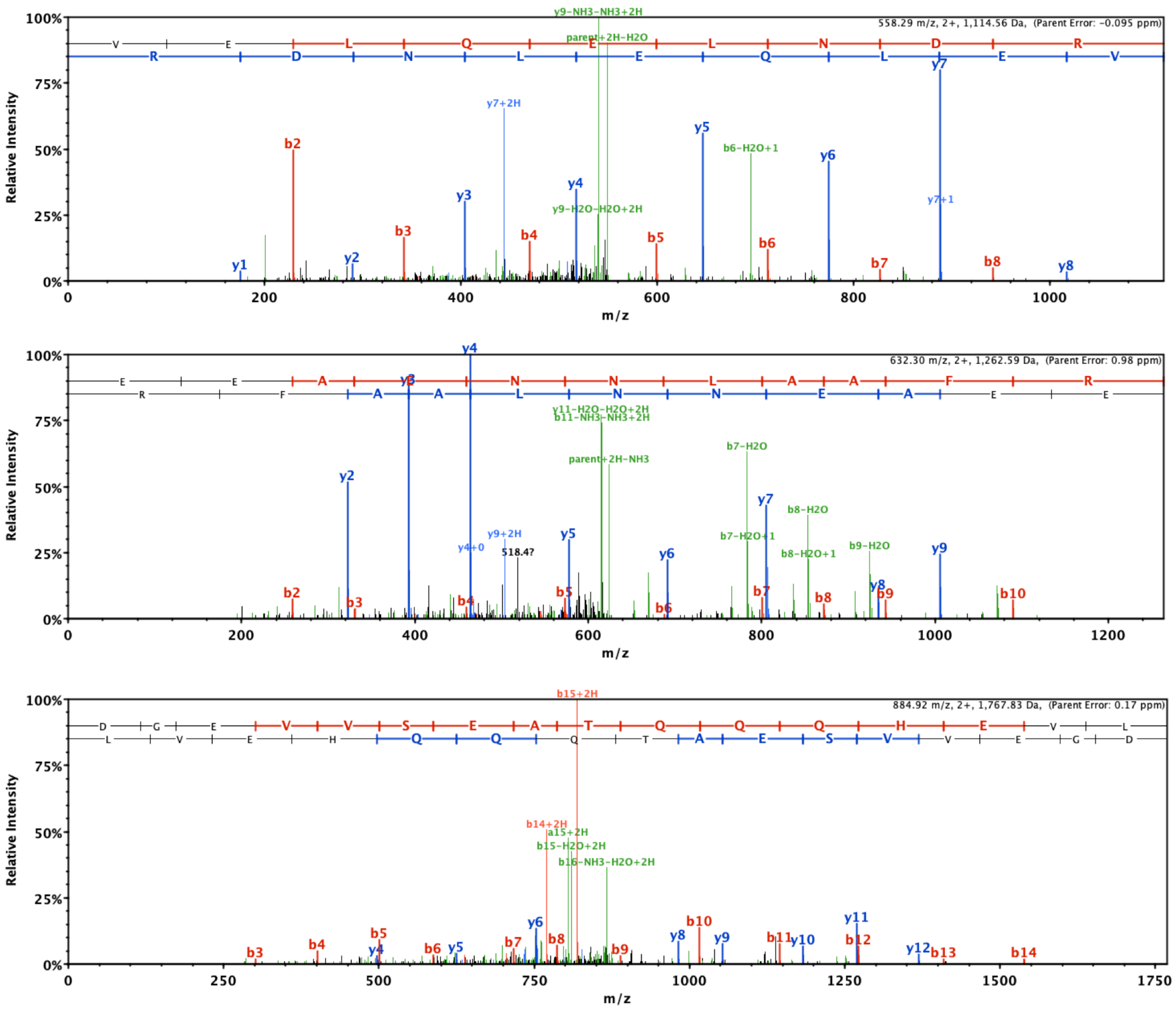


**c**

#### DES ~200 kDa, R120G cryAB *vs.* NTG

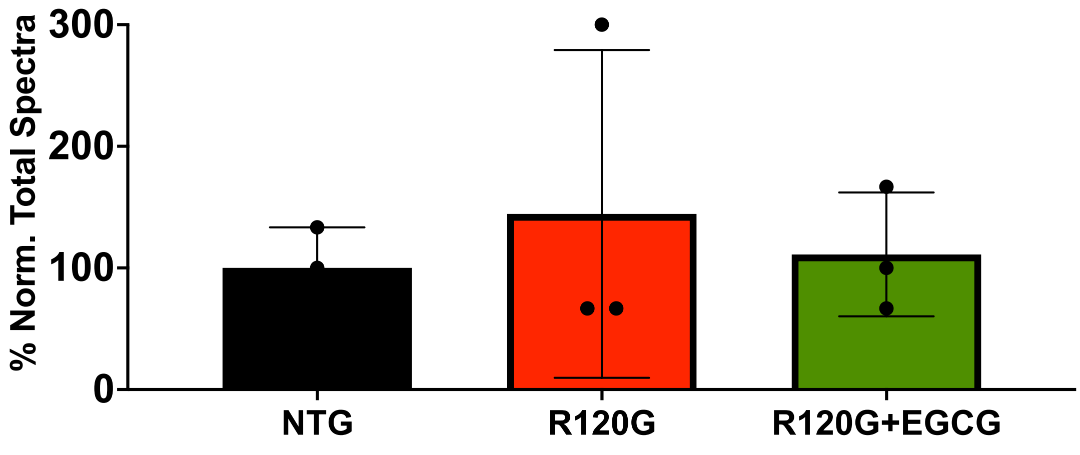

#### Supplementary Figure 2

**a**

TAU_HUMAN (100%), 78,927.7 Da
Microtubule-associated protein tau OS=Homo sapiens GN=MAPT PE=1 SV=5
2 exclusive unique peptides, 2 exclusive unique spectra, 4 total spectra, 28/758 amino acids (4% coverage)

```
MAEPRQEFEV MEDHAGTYGL GDRKDQGGYT MHQDQEGDTD AGLKESPLQT PTEDGSEEPG SETSDAKSTP
TAEDVTAPLV DEGAPGKQAA AQPHTEIPEG TTAEEAGIGD TPSLEDEAAG HVTQEPESGK VVQEGFLREP
GPPGLSHQLM SGMPGAPLLP EGPREATRQP SGTGPEDTEG GRHAPELLKH QLLGDLHQEG PPLKGAGGKE
RPGSKEEVDE DRDVDESSPQ DSPPSKASPA QDGRPPQTAA REATSIPGFP AEGAIPLPVD FLSKVSTEIP
ASEPDGPSVG RAKGQDAPLE FTFHVEITPN VQKEQAHSEE HLGRAAFPGA PGEGPEARGP SLGEDTKEAD
LPEPSEKQPA AAPRGKPVSR VPQLKARMVS KSKDGTGSDD KKAKTSTRSS AKTLKNRPCL SPKHPTPGSS
DPLIQPSSPA VCPEPPSSPK YVSSVTSRTG SSGAKEMKLK GADGKTKIAT PRGAAPPGQK GQANATRIPA
KTPPAPKTPP SSGEPPKSGD RSGYSSPGSP GTPGSRSRTP SLPTPPTREP KKVAVVRTPP KSPSSAKSRL
QTAPVPMPDL KNVKSKIGST ENLKHQPGGG KVQIINKKLD LSNVQSKCGS KDNIKHVPGG GSVQIVYKPV
DLSKVTSKCG SLGNIHHKPG GGQVEVKSEK LDFKDRVQSK IGSLDNITHV PGGGNKKIET HKLTFRENAK
AKTDHGAEIV YKSPVVSGDT SPRHLSNVSS TGSIDMVDSP QLATLADEVS ASLAKQGL
```

**b**

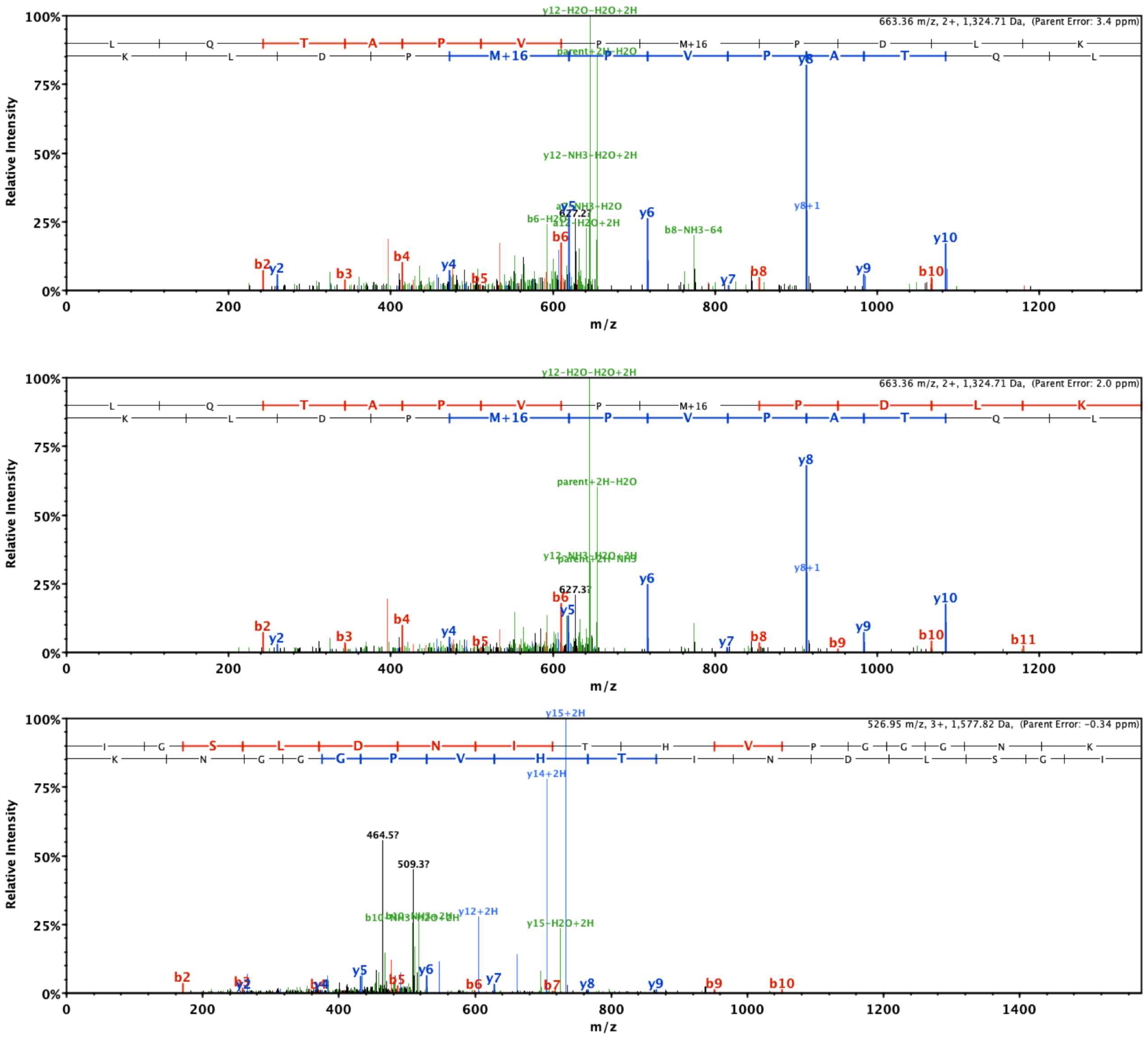


**c**

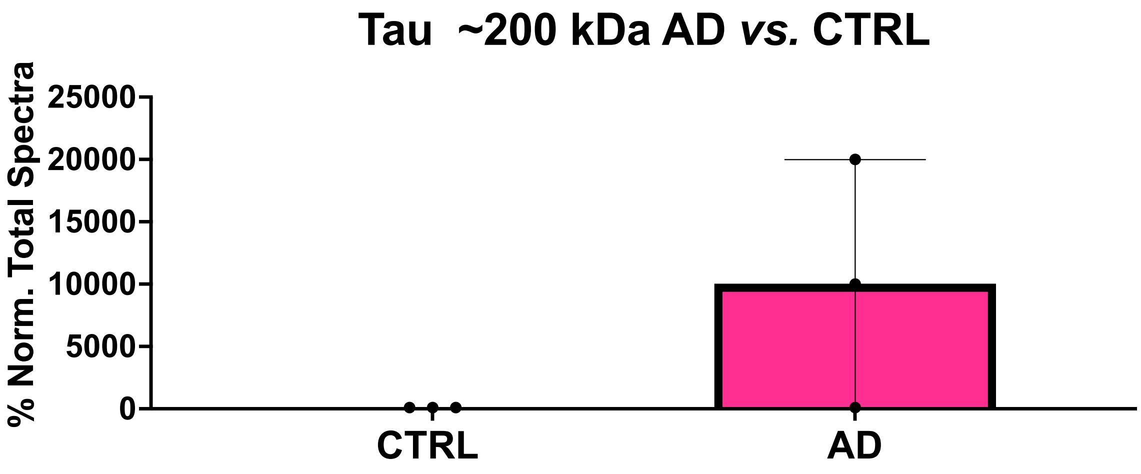
