## Supplemental Methods and Results for "Thioflavin T In-gel Stain to Study Protein Misfolding in Frozen Tissue Specimens"

**Supplementary Methods**

Mass spectrometry (MS) analysis: In-gel digestion was performed on bands which were located using Coomassie as a reference as previously described (Rainer PP et al., Circ Res 2018).

Liquid chromatography and tandem MS analysis (LC-MS/MS) were performed using an Ultimate 3000 nano LC (Thermo Scientific) connected to an Orbitrap Elite mass spectrometer (Thermo Scientific) equipped with an EasySpray ion source. Peptides were first loaded onto a trap column (PepMap100 C18, 5 $\mu$m, 100 Å, 300 $\mu$m i.d. x 5 mm, Thermo Scientific) followed by separation on a PepMap RSLC C18 column (2 $\mu$m, 100 Å, 75 $\mu$m i.d. x 25 mm, Thermo Scientific) using a flow rate of 300 nL/min with a linear gradient of 5-30%B for 30 minutes, 35-95%B for 3 minutes, holding at 95%B for 7 minutes and re-equilibrating at 5%B for 25 minutes at 400 nL/min (mobile phase A was 0.1% formic acid in water and mobile phase B was 0.1 % formic acid in acetonitrile). The nano-source capillary temperature was set to 275 °C and the spray voltage was set to 2 kV. MS1 scans were acquired in the Orbitrap Elite at a resolution of 60,000 FWHM (400-1700 m/z) with an AGC target of 1x106 ions over a maximum of 250 ms. MS2 spectra were acquired for the top 15 ions from each MS1 scan in CID mode in the ion trap and a target setting of 1x104 ions, an accumulation time of 100 ms, and an isolation width of 2 Da.

The normalized collision energy was set to 35% and one MicroScan was acquired for each spectrum. Monoisotopic precursor selection was enabled and only MS1 signals exceeding 500 counts triggered the MS2 scans, with +1 and unassigned charge states not being selected for MS2 analysis. Dynamic exclusion was enabled with a repeat count of 2, repeat duration of 30 seconds and exclusion duration of 90 seconds.

Tandem mass spectra were extracted by Proteome Discoverer (Thermo). All MS/MS samples were analyzed using Sequest (Thermo Fisher Scientific, San Jose, CA, USA; version 1.0). Sequest was set up to search the respective species-specific databases(mouse and human, uniprot) assuming the digestion enzyme with trypsin. Sequest was searched with a fragment ion mass tolerance of 1.00 Da and a parent ion tolerance of 50 PPM. Carbamidomethyl of cysteine was specified in Sequest as a fixed modification. Oxidation of methionine, phosphorylation of serine, threonine and tyrosine, glyGly of lysine and leuArgGlyGly of lysine were specified in Sequest as variable modifications.

Scaffold (version Scaffold_5.1.2, Proteome Software Inc., Portland, OR) was used to validate MS/MS based peptide and protein identifications. Peptide identifications were accepted if they could be established at greater than 95.0% probability by the Peptide Prophet algorithm (Keller, A et al., Anal. Chem. 2002) with Scaffold delta-mass correction. Protein identifications were accepted if they could be established at greater than 95.0% probability and contained at least 2 identified peptides. Protein probabilities were assigned by the Protein Prophet algorithm (Nesvizhskii, AI et al., Anal. Chem. 2003). Proteins that contained similar peptides and could not be differentiated based on MS/MS analysis alone were grouped to satisfy the principles of parsimony. Proteins sharing significant peptide evidence were grouped into clusters.

**Supplementary Results**

One of the advantages of the ThT in-gel protocol is its compatibility with downstream MS. Specifically, ThT-positive bands can be excised using Coomassie as reference and their composition can be analyzed by MS. First, we excised and analyzed by MS the ThT-positive bands at ~200 kDa from gels comparing cardiac protein extracts from R120G cryAB and NTG mice. We included R120G cryAB extracts treated with EGCG in the analysis (**Sup. Figure 1**). As we describe in detail in the main text this treatment reduces or reverses amyloid fibrillization *ex vivo*. Therefore, we used the EGCG-treated samples to strengthen the correlation between the increased abundance of desmin in ThT-bands with disease. When we subjected the peptides extracted from the ThT-positive band at ~200 kDa from NTG and R120G cryAB -/+ EGCG, we detected desimin, whose molecular weight is ~53 kDa (**Sup. Figure 1a-b**). This observation suggests that desmin tetramers could be involved in the formation of ThT-postive fibrils in this model. Further, a summary quantitation based on the normalized abundance of total, unique spectra for desmin peptides highlighted an increase in desmin levels in the R120G cryAB. The contribution of desmin to the increase in the ThT fluorescence was strengthen by the concurrent decrease of its peptides in R120G cryAB samples treated with EGCG (**Sup. Figure 1c**).

In addition, we subjected ~200 kDa ThT-positive gel bands from brain tissue protein extracts from AD patients and healthy controls (CTRL) to a similar analysis (**Sup. Figure 2**) and detected an increase in Tau in AD samples compared to controls (**Sup. Figure 2c**).

These data combined, suggest that the ThT in-gel platform could be coupled to targeted MS methods, such as parallel reaction monitoring (PRM) or data independent acquisition (DIA) (please see Agnetti G. *et al.*, Manual of Cardiovascular Proteomics, 2016 for details) to gain both sensitivity and quantitative accuracy.

**Legends to Supplementary Figures**

**Supplementary Figure 1. MS Analysis of ThT-positive Bands Confirms Desmin Involvement in Amyloid Fibrils Formation in the Heart of R120G cryAB Mice. a)** Representative sequence coverage for mouse desmin in ThT-positive bands at ~200 kDa in cardiac extracts from R120G cryAB mice. **b)** Representative spectra for desmin peptides in the same samples. **c)** Summary quantitation of normalized, total unique desmin spectra as provided by Scaffold 5.

**Supplementary Figure 2. Increased Tau Levles in ThT-positive bands from AD Brain Protein Extracts. a)** Representative sequence coverage for human Tau in ThT-positive bands at ~200 kDa from AD brain extracts. **b)** Representative spectra for Tau peptides in the same samples. **c)** Summary quantitation of normalized, total unique Tau spectra as provided by Scaffold 5.
